# A machine learning approach reveals that spore levels in organic bulk tank milk are dependent on farm characteristics and meteorological factors

**DOI:** 10.1101/2024.08.03.606478

**Authors:** Chenhao Qian, Renee T. Lee, Rachel L. Weachock, Martin Wiedmann, Nicole H. Martin

## Abstract

Bacterial spores in raw milk can lead to quality issues in milk and milk derived products. Since these spores originate from farm environments, it is important to understand contributions of farm-level factors to spore levels in raw milk. Identifying highly influential factors will guide interventions to control the transmission of spores from farm environments into bulk tank raw milk and therefore minimize spoilage in the finished products. The objective of this study was to investigate the impact of farm management practices and meteorological factors on levels of different spore types in organic raw milk by leveraging machine learning models. In this study, raw milk from certified organic dairy farms (n = 102) located across 11 states was collected 6 times over a year and tested for standard plate count, psychrotolerant spore count, mesophilic spore count, thermophilic spore count, and butyric acid bacteria. At each sampling date, a survey was collected from each farm to obtain structured data about farm management practices. Meteorological factors related to temperature, precipitation, solar radiation, and wind were obtained on the date of sampling as well as 1, 2, and 3 days prior to the date of sampling from an open-source website. The dataset was stratified separately based on the use of a parlor for milking, number of years since organic certification, and whether the lactating herd was exposed to pasture time into sub-datasets to address the potential confounders. Using the entire datasets and 6 sub-datasets respectively, we constructed random forest regression models to predict log_10_ mesophilic spore count, log_10_ thermophilic spore count, and log_10_ butyric acid bacteria most probable number as well as a random forest classification model to classify the presence of psychrotolerant spores in each raw milk sample. The summary statistics showed that spore levels vary considerably between certified organic farms but were only slightly higher than spore levels previously reported from conventional dairy farms. The variable importance plots from the random forest models suggest that herd size, certification year, employee-related variables (e.g., number of people milking cows per week), clipping and flaming udders, stocking density, and principal component representing air temperatures are among the top variables influencing the spore levels in organic raw milk, despite the limitation in the model performance (the highest performance for regression and classification is R^2^ of 0.36 for predicting TSC for farms with a parlor and accuracy of 0.73 for classifying positive PSC for farms without a parlor, respectively). The relatively small effects of top variables as demonstrated by the partial dependence plots suggest that an individualized approach that synergistically considers multiple farm and environmental factors is needed to enable a risk-based approach for managing spore levels. While at the current stage, these models were insufficiently accurate to be used as predictive tools, incorporating novel data streams such as video surveillance and daily farm observations with computer vision and natural language processing, respectively, has the potential to enhance the performance of the model as a real-time monitoring tool for spores as an indicator of milk microbiological quality.

## INTRODUCTION

Maintaining raw milk microbial quality is crucial for ensuring the safety and quality of finished dairy products. Sporeforming bacteria, which represent a diverse group of microorganisms that have various optimal growth conditions and spoilage mechanisms, are a key quality parameter for raw milk. Due to the biological diversity, different groups of sporeforming bacteria have the potential to induce a variety of spoilage defects in a range of finished products, including pasteurized milk and semi-hard cheese, as well as beverages made with dry dairy ingredients (Martin et al., 2021, 2023). Major groups of sporeforming bacteria that have been previously associated with dairy spoilage include (i) mesophilic (e.g., *Bacillus licheniformis*) and thermophilic sporeformers (e.g., *Anoxybacillus flavithermus*, *Geobacillus* spp.), which can survive and grow at relatively high optimal temperatures (approximately 32 and 55 C°, respectively) during the powder manufacturing process, leading to formation of biofilms that could contaminate powder during processing (Rückert et al., 2004; Delaunay et al., 2021); (ii) psychrotolerant sporeformers (e.g., *Paenibacillus* spp.), which can grow at low temperatures (e.g., 6 C°) typical for cold storage and transportation of high-temperature, short-time pasteurized fluid milk and therefore can cause milk to develop defects (e.g., off-flavor and coagulation) at the late stage of shelf life (Fromm and Boor, 2004; Huck et al., 2007); and (iii) anaerobic butyric acid bacteria (e.g., *Clostridium tyrobutyricum*), some of which have high salt tolerance and therefore can grow in the brined semi-hard cheese at the late stage of ripening and induce late blowing defects characterized by gas formation and acidic off-flavor development (Garde et al., 2013; D’Amico, 2014; Düsterhöft et al., 2017). These diverse groups of sporeforming bacteria pose various challenges to maintaining high-quality dairy products throughout the shelf-life because of their ability to survive pasteurization and grow under the favorable conditions of shelf-life storage. Therefore, even at very low initial levels, these sporeforming bacteria can still pose a concern for the quality of finished product.

While it is feasible to extend the shelf life and improve the quality of dairy products by reducing the level of sporeforming bacteria at the processing level through technological solutions, such as centrifugation and bactofugation (Hoffmann et al., 2006; Schmidt et al., 2012; Doll et al., 2017), controlling the contamination at the farm level by improving farm practices is equally important since the farm environment represents a major entry point for these sporeforming bacteria. A number of studies (Masiello et al., 2014, 2017; Miller et al., 2015; Martin et al., 2019; Murphy et al., 2019; Evanowski et al., 2020, 2023) have reported the association of farm management practices with different types of sporeformers in raw milk. For instance, practices that aim to improve udder hygiene, such as clipping and flaming udder hair (Martin et al., 2019) and fore stripping (Miller et al., 2015; Murphy et al., 2019), were shown to be associated with bulk tank spore levels. In addition, bedding materials were also reported in several studies (Magnusson et al., 2007; Murphy et al., 2019) to influence the spore levels in raw milk. However, all of these studies were focused on the conventional farming system. Because organic dairy farms have different management requirements than conventional farms, the implications for different practices and environmental conditions on the spore levels may differ. For example, organic dairy farms need to provide pasture access for cows no fewer than 120 days a year, which might expose cows to more sporeformers in the natural environment which is considered a primary source for these organisms.

Therefore, the objective of this study is to investigate prevalence and levels of spores in organic raw milk as well as understanding the factors that might be correlated with these spores. We proposed using a machine learning method to analyze high-dimensional farm-level data collected via intensive samplings from 102 organic farms across multiple states in a one-year time frame. Machine learning methods have been increasingly used to understand practices or environmental conditions that can drive food safety and quality issues. In one study (Murphy et al., 2021), a conditional random forest was constructed to understand the relative importance of various management factors (e.g., annual training, type of cleaning-in-place) in a dairy processing facility. For a different food domain (i.e., fresh produce) but with similar concepts, numerous studies (Weller et al., 2020, 2021; Green et al., 2021) have attempted to predict and investigate pathogen levels in agricultural water for fresh produce using publicly available databases that contain meteorological and geospatial data. These meteorological effects could be relevant to dairy farms as well considering that cows in organic dairy farms spend time on pasture where spores are prevalent, and we hypothesize that the meteorological conditions are likely influencing the transmission of spores from pasture environments (e.g., soil) to the udder. Therefore, we believe that a machine learning approach will be appropriate for the type of data and tasks we have in our study. Ultimately, this study provides the organic dairy industry with key factors associated with the presence and levels of dairy relevant groups of spores. Future studies can leverage these results to design practical interventions for reducing bacterial spores in organic bulk tank milk.

## MATERIALS AND METHODS

### Sample Collection and Microbiological Testing

Raw milk samples (∼295 mL/sample) from certified organic dairy farms (n = 102) in 11 US states (Table 1) were collected by milk truck drivers and project staff in sterile vials. Samples were collected bimonthly between May 2021 and Aug 2022 (for a total of 6 samples per farm) from the top of the bulk tank after 5 minutes of agitation on each farm using a sanitized dipper as previously described (Evanowski et al., 2020). The samples were then frozen and shipped on ice to the Cornell Milk Quality Improvement Program (MQIP; Ithaca, NY). Upon arrival, the samples were kept frozen at -20° for up to 30 days until thawed for analysis at 6°C for 24h prior to conducting microbiological testing for mesophilic spore count (MSC), thermophilic spore count (TSC), psychrotolerant spore count (PSC) and butyric acid bacteria (BAB). Milk samples were heat treated (HT) at 80°C for 12 mins (Frank and Yousef, 2004). Then 1 mL was pour plated in duplicate using standard methods agar (SMA) and incubated at 32°C and 55°C for MSC and TSC, respectively. For PSC, a 15-tube most probable number (MPN) technique was used with 5 tubes each of the following prepared for each raw milk sample: (i) 10 mL of HT milk, (ii) 1 mL of HT milk in 10 mL skim milk broth (SMB), and (iii) 0.1 mL of HT milk in 10 mL SMB. Tubes were incubated at 6°C for 21 days; then each tube was evaluated for growth by streaking 10 µL of sample onto SMA with a sterile loop prior to incubation at 32°C for 48h. To test for BAB, raw milk samples were transferred into 9 mL Bryant and Burkey (BB) broth using a 20-tube MPN scheme of (i) 10 tubes of 5 mL and (ii) 10 tubes of 0.5 mL. Molten paraffin was used to create an anaerobic environment in the BAB MPN tubes by covering the media and milk with 2 cm of wax. Samples were subsequently heat treated at 75°C for 15 mins to eliminate any remaining vegetative cells before incubation at 37°C for 6 days. Gas production in tubes was assessed every 48 hours. Any tubes with observed movement of the wax plug, indicating gas formation, were scored as positive. The number of positive and negative tubes for each sample was used to calculate the final BAB MPN/mL as previously described (Shi et al., 2023).

**Table 1.**
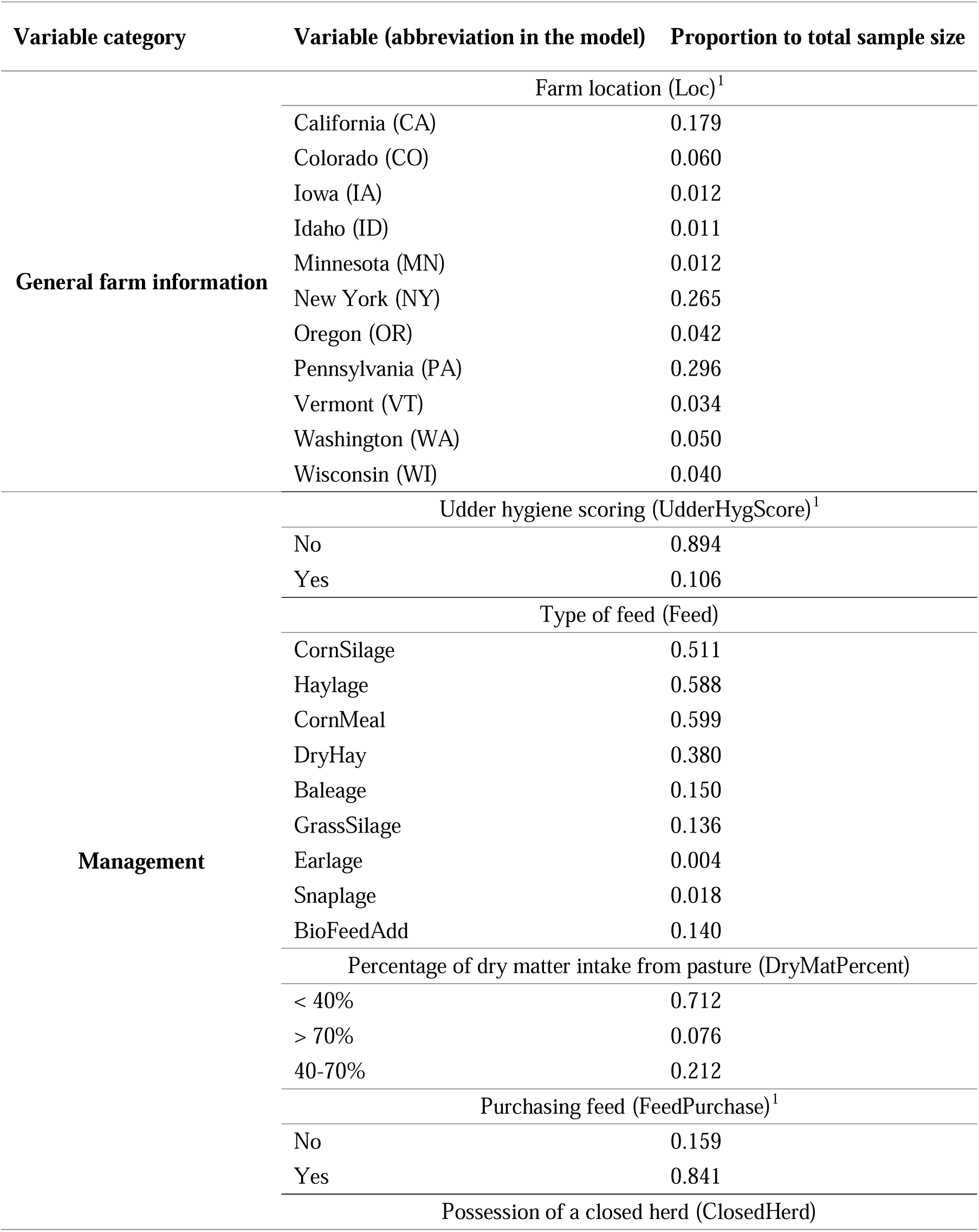
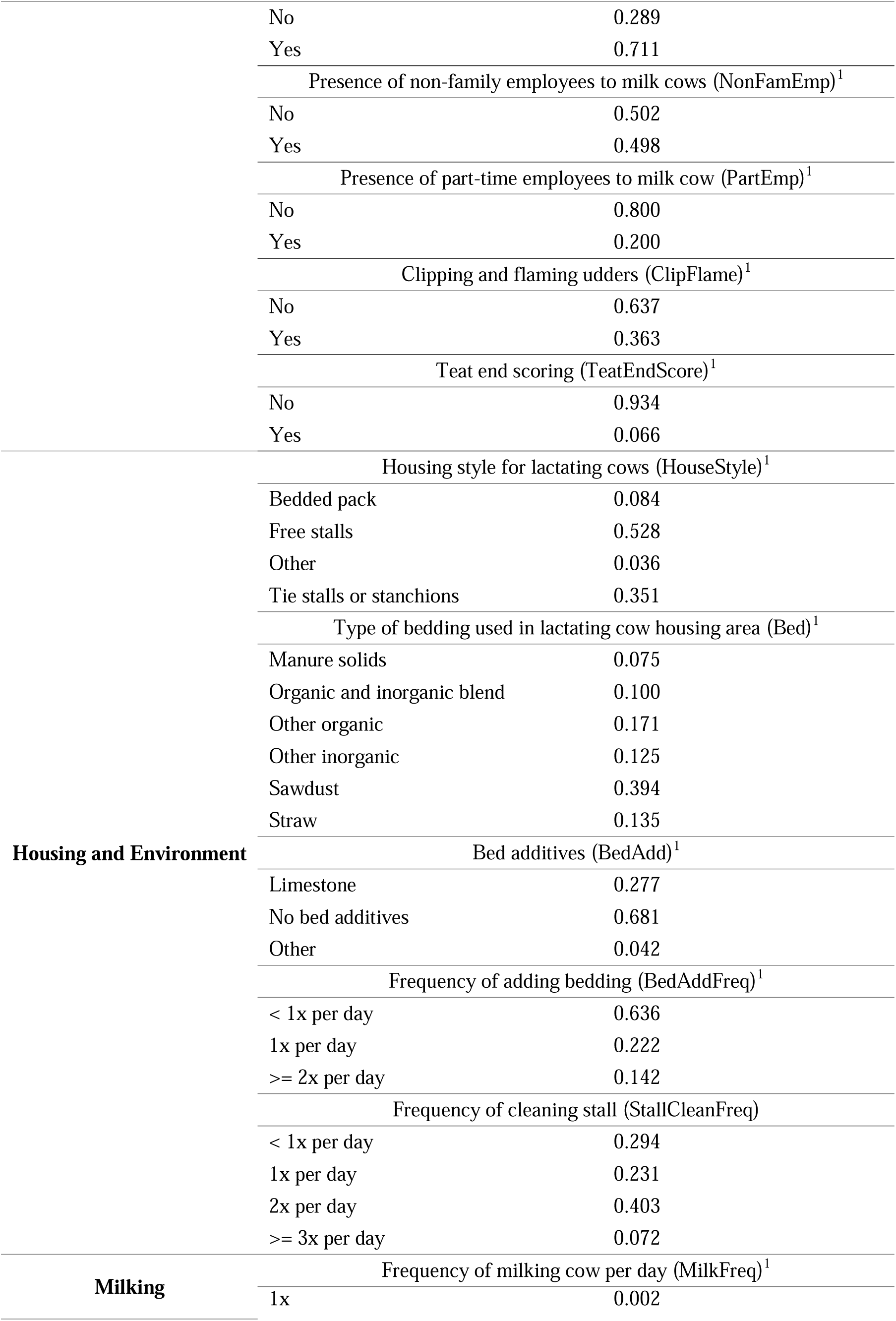
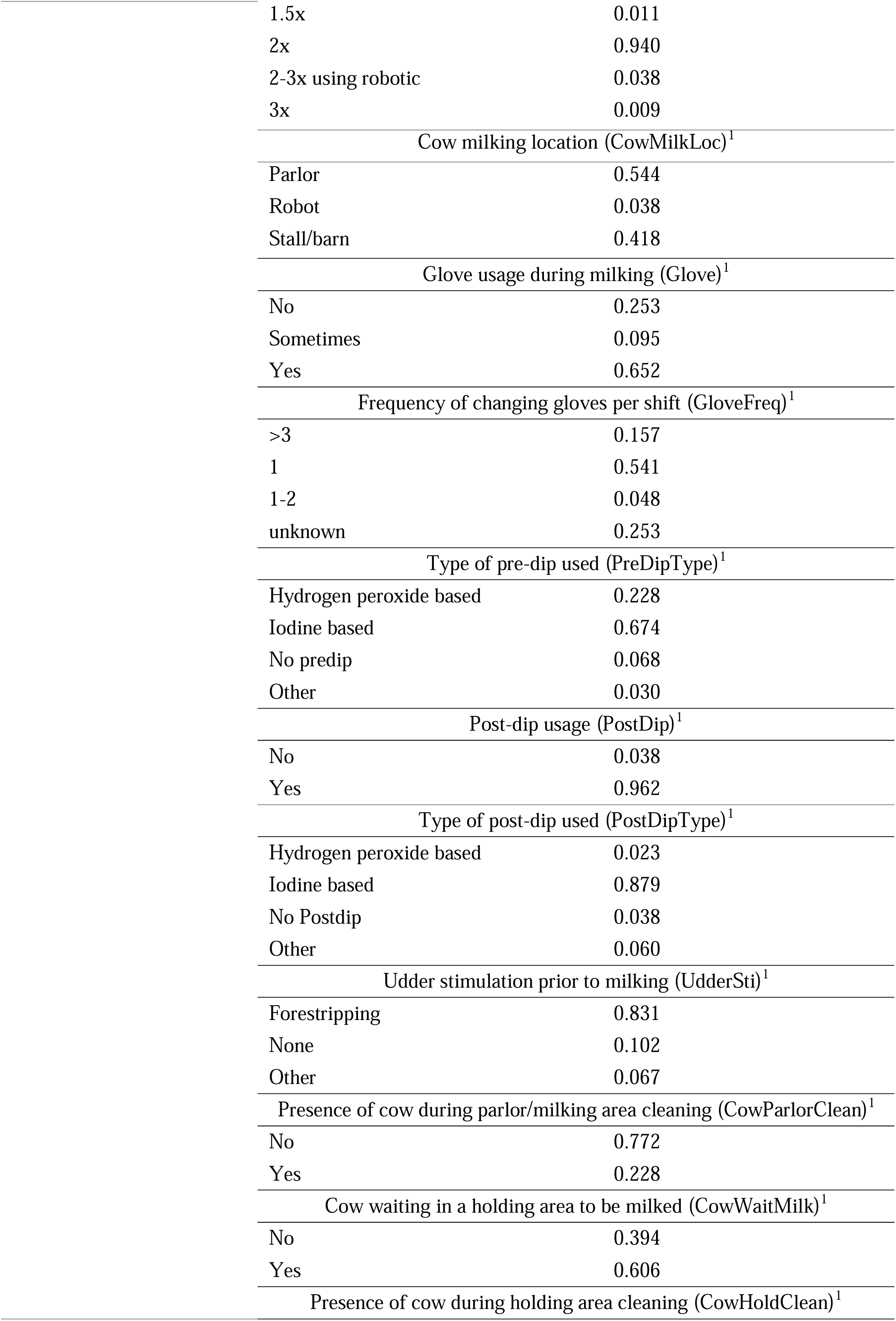
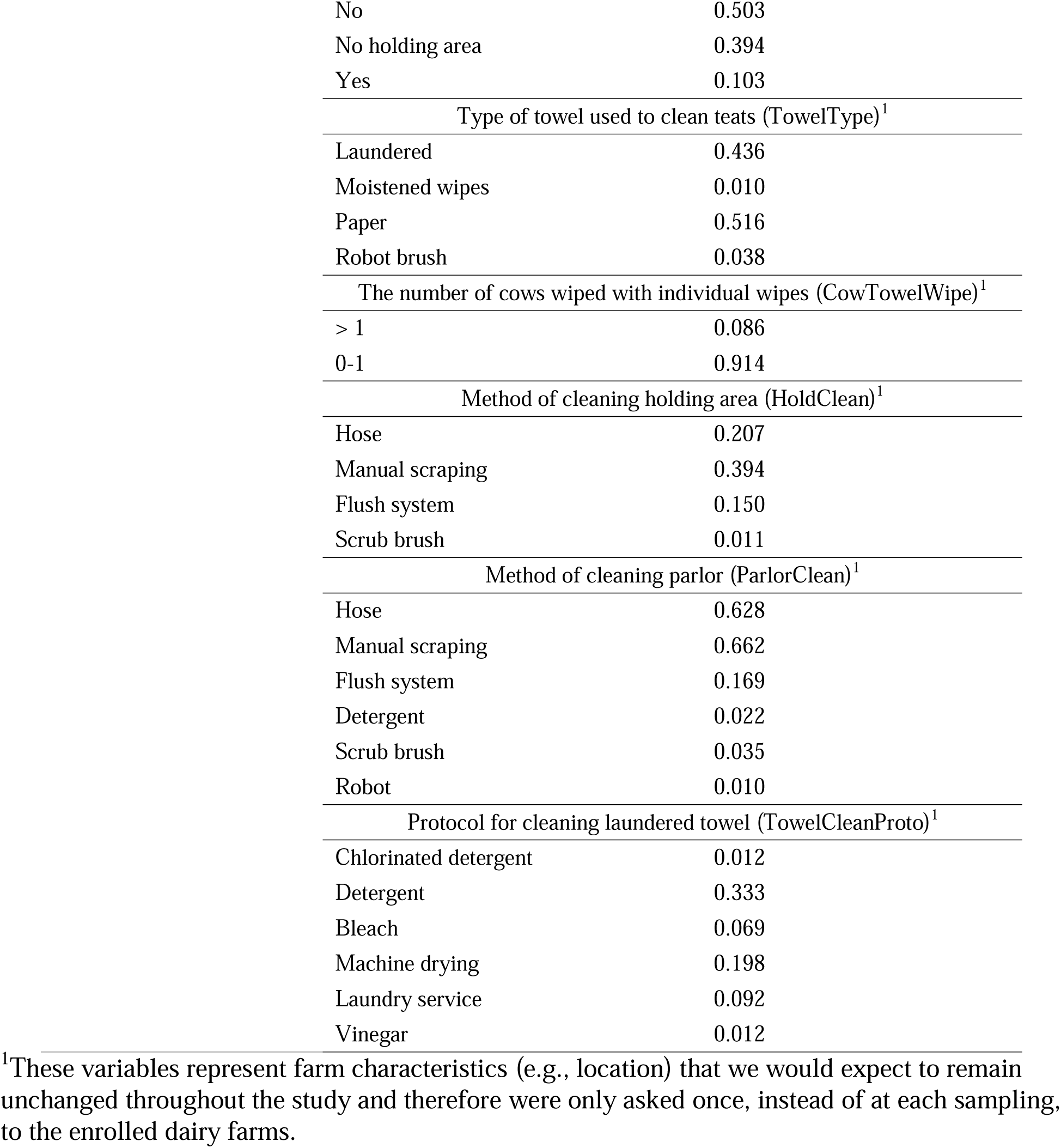
Summary statistics of preprocessed categorical variables collected in a survey administered to 102 certified organic dairy producers. Total proportions of non-exclusive answers for multiple choice questions (e.g., type of feed) can add up to more than 1. The abbreviations for each variable are presented in parentheses.

In addition to spore testing, we also performed standard plate count (SPC) to assess the total bacterial levels in collected organic raw milk samples. SPC testing was conducted in duplicate using a spiral plater (Neu-tec Eddy Jet 2, Farmingdale, NY). We plated 50 μL of undiluted (10^0^) and diluted (10^-2^) milk samples onto standard methods agar (EMD Millipore Corporation, Billerica, MA), and then incubated the plates for 48 hours at 32°C prior to enumeration using an automated colony counter (SphereFlash; Neu-tec Automated Colony Counter, Farmingdale, NY) (Laird et al., 2004).

### Survey Development and Administration

A farm management practices survey was administered at the time of each sample collection on each farm. The survey was developed by the Milk Quality Improvement Program at Cornell University. For the first round of sampling, survey responses were collected over the phone or in person by project staff from Cornell University, University of California Davis, Oregon State University, Penn State University, and University of Nebraska-Lincoln, and survey responses were provided by farm owners or herd managers. For second and subsequent samplings, the survey was administered by project staff over the phone, in person, over email, or through the mail to accommodate the preferences of each farmer. The survey consisted of multiple questions (see https://github.com/FSL-MQIP/OREI_ML_Inference/tree/main/Data for questions and answers from the raw survey data) in four major sections, including general farm info (e.g., number of cows), management (e.g., udder clipping and flaming), housing and environment (e.g., stall cleanliness), and milking (e.g., udder preparation method).

All survey responses were submitted by project staff at collaborating universities to an online survey tool (Qualtrics, Provo, UT) for analysis to be conducted by project staff at the Milk Quality Improvement Program at Cornell University.

### Descriptive Analysis

Histograms were created for SPC and each spore count across all collected samples to understand the overall variability in different types of microbial indicators (i.e., SPC and spore count). In addition, the mean and standard deviation were calculated separately for SPC and each type of spore count from samples of each individual farm. The distribution of the mean was used to indicate the between-farm variability while the distribution of the standard deviation was used to indicate within-farm variability. The purpose of this descriptive analysis is to differentiate the source of variation in the SPC as well as spore count.

### Modeling

#### Data Collection and Preprocessing

Raw survey data were first processed prior to inclusion as predictor variables. The identifiers for farms, such as state and co-op were excluded from the list of predictor variables, as these identification factors are highly confounded (i.e., reflecting many intertwined factors such as climate and employee demographics) and therefore interpreting their effects could potentially be misleading. Survey responses were further examined and processed to combine any responses that had extremely low frequency as some responses were only provided by one farm and therefore not representative. For example, bedding materials including ashes, limestone, gypsum, and oyster shells were grouped as “other”.

Meteorological factors on the sampling date as well as 1-3 days prior to the sampling date were collected from an online API Visual Crossing (https://www.visualcrossing.com/). The meteorological data at the nearest weather station to the farm sampled was used to approximate meteorological factors at that farm. Collected variables include daily maximum temperature, minimum temperature, average temperature, humidity, precipitation, precipitation cover, wind gust, wind speed, and solar radiation.

In addition to the survey and meteorological variables, we included a new variable, the variability (i.e., standard deviation) of SPC for each farm as previously described, in the list of predictor variables. This variable was used as a proxy for the ability of a farm to maintain the bacterial levels in their raw milk (e.g., farm with lower SPC variability might be more proficient in quality assurance). If a farm has only one sample tested for SPC, then the SPC variability was imputed using the mean SPC variability across all farms.

All predictor variables were preprocessed to improve training efficiency and eliminate weak predictors that are sparse and might bring only marginal benefit to the model. First, all numeric predictors were normalized via centering and scaling and all nominal predictors were converted into dummy variables. Then, due to high dimension and collinearity, using principal component extractions, meteorological factors were represented by the top five principal components that explained the most variability within the data. Lastly, variables that have highly imbalanced responses were removed using near-zero variance filter with the frequency distribution ratio set to 85/15.

#### Model Setup

Regression was performed separately with log_10_MSC, log_10_TSC, and log_10_BAB as the numeric outcome variables while classification was performed with MPN results of PSC as the binary outcome variable. The decision to develop a classification model only for PSC was because psychrotolerant sporeformers were detectable in only 72.1% raw milk samples and therefore developing a regression model for zero-inflated outcome variable might be biased. Models were independently trained using 7 data subsets, which consist of 1 dataset containing all available data, 2 datasets stratified based on whether the farm uses a parlor for milking cows or an alternative approach (e.g., milking in tie stalls), 2 datasets stratified based on the number of years the farm has been certified organic (i.e., whether the farm has been certified for ≤ 9 years or > 9 years), and 2 datasets stratified based on, at each sampling date, whether the lactating cows were exposed to pasture or not. While stratification inevitably reduced the sample size, it can remove the potential confounding variables (Dohoo et al., 2003a) and allows more targeted suggestions about practices to specific farm management styles. In addition, stratification can create data subsets with more homogenous characteristics and therefore potentially improve the model performance.

#### Random Forest Analysis

The random forest models were developed using tidymodels (Kuhn and Wickham, 2020) in R programming language (R Core Team, 2021).The tree-based method was selected as this approach can handle high-dimensional categorical variables and is particularly suitable for strong predictors (Borup et al., 2023), which we assume farm management practices and meteorological factors to be. Each model was trained by 3-fold cross-validation repeated 5 times to tune the hyperparameter “min_n” and “mtry”, which stand for the minimum number of samples required to split the nodes, and the number of variables to sample at each split, respectively. The tuning process was efficiently conducted using grid search of 40 combinations of hyperparameters to explore the space of possible hyperparameter configurations. In terms of evaluation metrics for the model, the coefficient of determination (R^2^) was used for each regression model while the accuracy was used for classification model. For classification models that showed imbalanced distribution of outcome (i.e., presence of PSC), oversampling at 1:1 ratio was performed using Synthetic Minority Oversampling Technique (SMOTE) to prevent the bias towards the outcome with higher proportion in the dataset (Chawla et al., 2002). To understand the relative importance of predictor variables to the model outcome, variable importance plots (VIPs) based on impurity reduction were constructed for each model developed (for a total of 28 VIPs). The top 10 variables for each VIP were retained and summarized in a heatmap sorting variables from highest frequency of appearance to the least. To further identify the directionality of the effect of predictor variables, we constructed partial dependence plots (PDPs) that show the expected change in spore levels with respect to the farm-related variables identified from VIPs. However, these PDPs are only intended to identify associations, rather than infer any causal relationships.

#### Code availability

The R codes for descriptive analysis and modeling are available on GitHub (https://github.com/FSL-MQIP/OREI_ML_Inference/blob/main/Data%20analysis.Rmd).

## RESULTS

### Bulk tank spore levels and farm characteristics vary considerably between certified organic farms

Across all 534 samples collected, the overall mean and standard deviation for SPC were 3.18 log_10_CFU/mL and 0.66 log_10_CFU/mL (Figure 1), respectively. To further investigate the variability between farms, we summarized the mean and standard deviation of SPC by each individual farm and calculated the summary statistics for these two values across all farms (Figure 2). The results showed that the mean SPC (which can be used to represent between-farm variability) by each individual farm ranged from 1.88 to 4.65 log_10_CFU/mL, with a mean of 3.19 log_10_CFU/mL and a standard deviation of 0.47 log_10_CFU/mL while the SPC standard deviation (which can be used to represent within-farm variability) by each individual farm with more than one sample tested for SPC ranged from 0.06 to 2.29 log_10_CFU/mL with a mean of 0.45 log_10_CFU/mL and a standard deviation of 0.36 log_10_CFU/mL. These results indicate that within-farm variability differs considerably between farms as the standard deviation (i.e., 0.36 log_10_CFU/mL) is similar to the mean (i.e., 0.45 log_10_CFU/mL), which implies a wide and flat distribution. The variability of SPC standard deviation between farms suggests that farms exhibit different degrees of control for bacterial levels in the raw milk. However, less variability does not necessarily equal higher microbial quality of milk, as raw milk can have consistently inferior quality. For example, the farm in our study with the lowest SPC standard deviation (i.e.,0.06 log_10_CFU/mL) showed a relatively high mean SPC (i.e., 3.04 log_10_CFU/mL) while the farm in our study that had the lowest mean SPC (i.e., 1.88 log_10_CFU/mL) had a relatively large SPC standard deviation (i.e., 0.32 log_10_CFU/mL).

**Figure 1.**
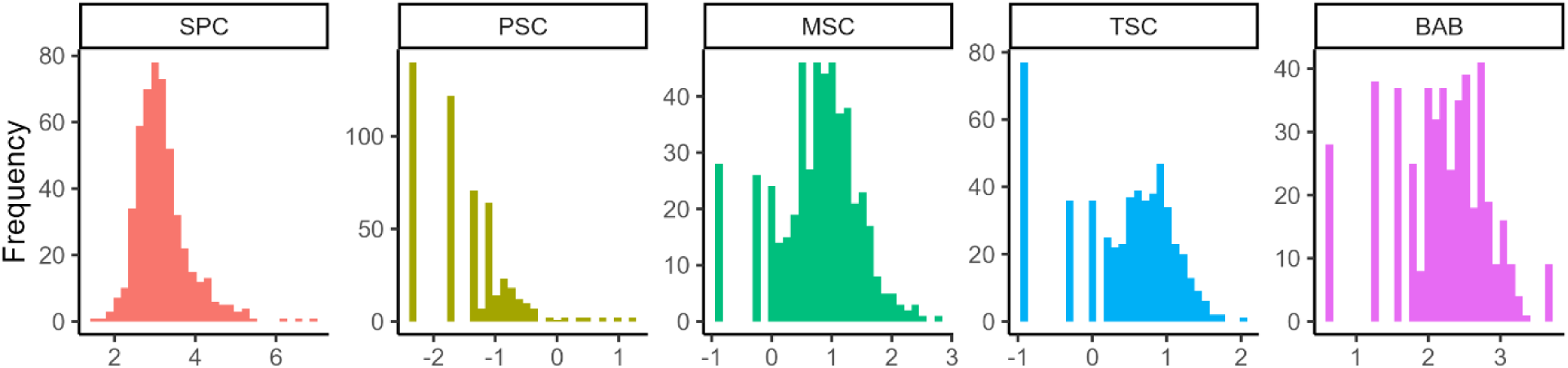
The distribution of overall standard plate count (SPC in log_10_CFU/mL; n = 534), psychrotolerant spore count (PSC in log_10_MPN/mL; n =534), mesophilic spore count (MSC in log_10_CFU/mL; n = 525), thermophilic spore count (TSC in log_10_CFU/mL; n = 556), and butyric acid bacteria (BAB in log_10_MPN/L; n = 497) across all samples collected from all farms (n = 102) enrolled in this study. The leftmost bar for each microbiological test plot represents an imputed value of 25% of detection limit after log_10_ transformation, which was used for all left-censored data as it is assumed that bacteria are still present at low levels below the detection limit.

**Figure 2.**
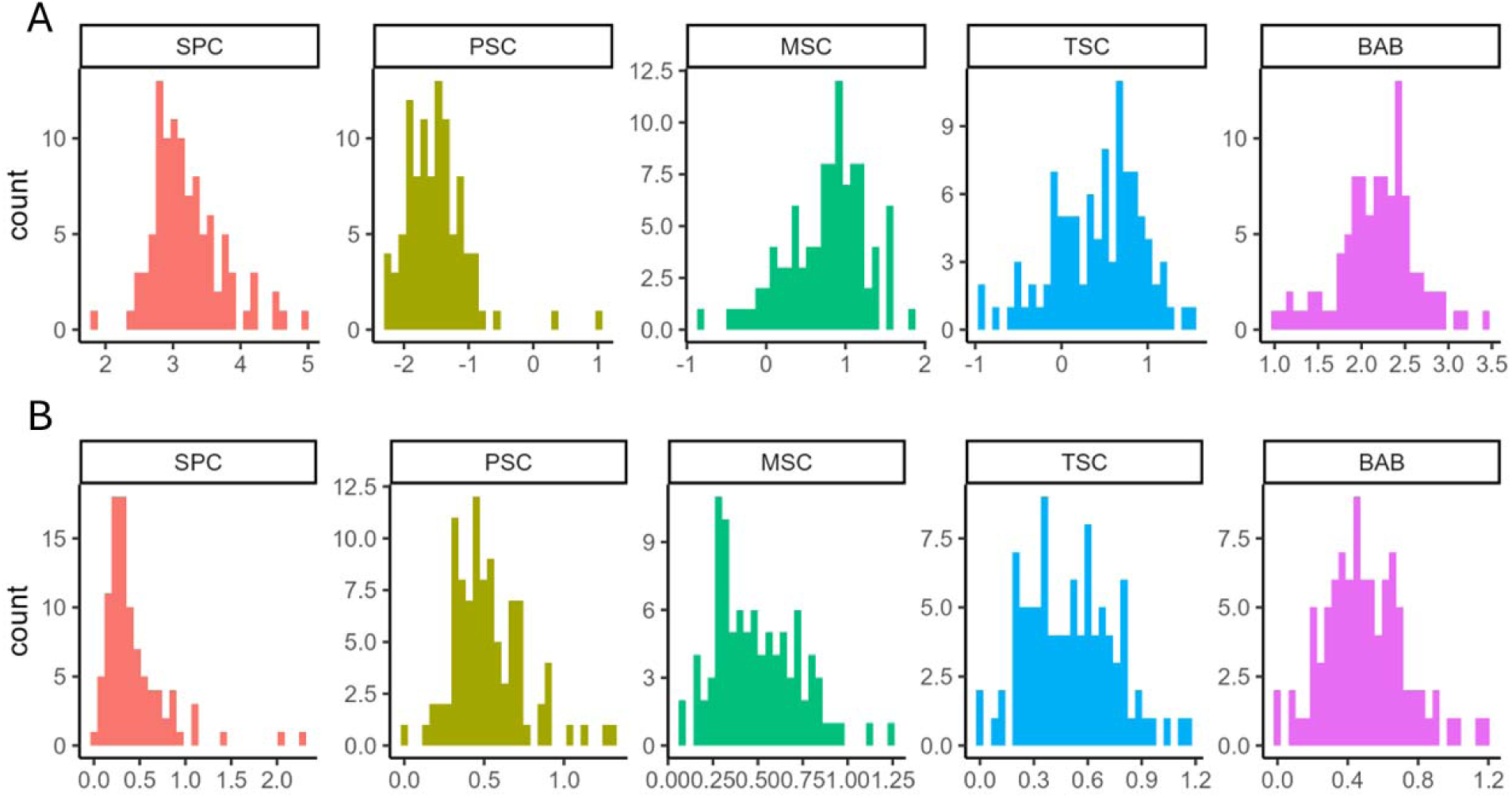
The distribution of mean (panel A) and standard deviation (panel B) for standard plate count (SPC in log_10_CFU/mL), psychrotolerant spore count (PSC in log_10_MPN/mL), mesophilic spore count (MSC in log_10_CFU/mL), thermophilic spore count (TSC in log_10_CFU/mL), and butyric acid bacteria (BAB in log_10_MPN/L) calculated for each individual farm enrolled in this study. (In the same order, coefficients of variance for the mean are 0.15, -0.30, 0.70, 1.27, and 0.21, and the standard deviations are 0.80, 0.43, 0.45, 0.48, and 0.46, respectively)

Similarly, we conducted summary statistics (Figure 1 and 2) for 4 different spore types using the overall data as well as summarized data by each individual farm. For each spore type, left censored data (i.e., samples where there were no detectable colonies) was replaced with a value equal to 25% of the detection limit for each test type (i.e., 0.5 CFU/mL for MSC, 0.5 CFU/mL for TSC, 20 MPN/L for PSC, and 18 MPN/L for BAB) prior to determination of mean and standard deviation. In a total of 525 samples collected for MSC, 94.3% (495/525) samples had MSC above detection threshold. The overall concentration of mesophilic spores (i.e., log_10_MSC) was normally distributed (Figure 1), with an overall mean of 0.76 log_10_CFU/mL and a standard deviation of 0.68 log_10_CFU/mL. The prevalence of BAB was found to be similar to MSC. In a total of 497 samples analyzed for BAB, 94.4% (469/497) samples had BAB above the detection threshold. The log_10_BAB results were normally distributed (Figure 1), with a mean of 2.19 log_10_MPN/L and a standard deviation of 0.65 log_10_MPN/L. Compared to both MSC and BAB, TSC results showed fewer samples above the detection limit (85.6%; 476 out of 556 samples) while the concentration distribution for all samples showed a normal distribution (Figure 1), with a mean of 0.43 log_10_CFU/mL and a standard deviation of 0.70 log_10_CFU/mL. Among 4 spore types tested, psychrotolerant sporeformers showed the lowest prevalence, with only 72.1% (385 out of 534) samples testing positive, and in total 97.2% (520 out of 534) samples containing less than 1 spore per mL. The mean and standard deviation for psychrotolerant sporeformers were -1.50 and 0.68 log_10_MPN/mL, respectively. In addition to the overall variability, we calculated between-farm and within-farm variability in spore levels as previously described for SPC. Using the coefficient of variance (CV) calculated for each distribution (see Figure 2), we found that generally SPC showed less between-farm variability (CV for mean SPC presented in Figure 2A is 0.15) and more within-farm variability (CV for SPC standard deviation presented in Figure 2B is 0.80, nearly twice of that for spores) compared to spores, suggesting that spore levels differ more by farm but yet differ less between samples collected at each farm. This finding can be attributable to the relatively low levels of spores found in milk compared to SPC.

The high variability in total bacterial count and spore levels are potentially attributable to the heterogeneous characteristics in the enrolled farms that are managed differently. Farms in this study were recruited from 11 states (Table 1) and represented a range of experiences as certified organic dairy producers ranging from 1 to 37 years of certification (Table 2). Farms ranged in size from 24 to 4000 cows, with a mean and standard deviation of 390 and 764 cows, showing considerable variability in farm size. Similarly, this high variability in farm size was observed in the number of non-family, full-time, and part-time employees milking cows, as these numbers all had higher standard deviation than the mean (Table 2). In terms of milking and housing styles, the majority (54.4%) of farms milked cows in a parlor, while the remaining farms utilized tie stalls (41.8%) and a smaller number utilized robotic milking systems (0.04%). Farms utilizing free stall housing systems accounted for 52.8% of recruited farms, while tie stalls, bedded pack, and other accounted for 35.1, 8.4, and 3.6% respectively. As our surveys were conducted across more than one year, the reported pasture time ranged from 0 to 24 hours, with a mean and standard deviation of 8.55 and 7.69 hours. At the time of survey administration, the dry matter intake from pasture was less than 40% (71.2% of the sampling points), between 40 and 70% (7.6% of the sampling points) and more than 70% (21.2% of the sampling points). When cows were not relying on the dry matter intake from the pasture, diverse types of feed were utilized, with corn silage (51.1%), haylage (58.8%), corn meal (59.9%), and dry hay (38.0%) as top choices. While most farm management practices and choices are heterogeneous, for certain management practices, we observed more homogeneity. For example, 94.0% of farms milk cows twice a day, 93.4% of farms did not score teat end cleanliness, 89.4% of farms did not score udder hygiene, 96.2% of farms applied post-dip after milking, and 91.4% of farms did not wipe more than one cow with each individual wipe or towel (Table 1).

**Table 2.** Summary statistics of preprocessed numeric variables collected in the survey administered to 102 certified organic dairy producers. The abbreviations for each variable are presented in parentheses.

| Variable category | Variable (abbreviation in the model) | Mean (±standard deviation) | Range |
| --- | --- | --- | --- |
| **General farm info** | Certification year (CertYear)<sup>1</sup> | 13.79 ± 6.63 | [1, 37] |
| | Number of cows (CowNum) | 390 ± 764 | [24, 4000] |
| **Management** | Pasture time in hours per day (PastureTime) | 8.55 ± 7.69 | [0, 24] |
| | Stocking density (StockDen) | 0.94 ± 0.54 | [0.00, 3.64] |
| | Number of different people milking cows per week (PplNumPerWk)<sup>1</sup> | 4.15 ± 3.82 | [0, 24] |
| | Number of people milking cows in one milking shift (PplNumPerShift)<sup>1</sup> | 1.84 ± 0.95 | [0, 6] |
| | Number of non-family employee milking cows (NonFamEmpNum)<sup>1</sup> | 2.36 ± 4.18 | [0, 24] |
| | Number of full-time employee milking cows (FullEmpNum)<sup>1</sup> | 2.36 ± 4.10 | [0, 24] |
| | Number of part-time employee milking cows (PartEmpNum)<sup>1</sup> | 0.40 ± 1.14 | [0, 8] |

<sup>1</sup>These variables represent farm characteristics that we would expect to remain unchanged throughout the study and therefore were only asked once, instead of at each sampling, to the enrolled dairy farms.

### Cow numbers, certification year, SPC variability, and employee-related factors show importance for organic raw milk spore levels, although there are limitations with model performance

We constructed variable importance plots for a total of 28 random forest models (as combinations of 4 spore types and 7 datasets including i) overall dataset, ii) data from farms with parlors, iii) data from farms without parlors, iv) data from farms when cows are on pasture, v) data from farms when cows are not on pasture, vi) data from farms that have been certified organic for at most 9 years, and vii) data from farms that have been certified organic for more than 9 years), and these plots are publicly available at https://github.com/FSL-MQIP/OREI_ML_Inference/tree/main/Figure/VIP. We further summarize the variables considered important in these VIPs in a heatmap, ranking these variables in descending order by the number of appearances across the 28 total models as a proxy for the order of relative importance (see Figure 3). To further illustrate the effect of variables identified important in VIP, we constructed PDPs (publicly available at https://github.com/FSL-MQIP/OREI_ML_Inference/tree/main/Figure/PDP) to estimate, on average, the change in spore levels (in the case MSC, TSC, and BAB) or likelihood of spore presence (in the case of PSC) with respect to those variables, to supplement the findings from VIPs. While the degree of change in the outcome was small in these PDPs compared to the variability of spore levels, we still observed some general trends for enrolled farms.

**Figure 3.**
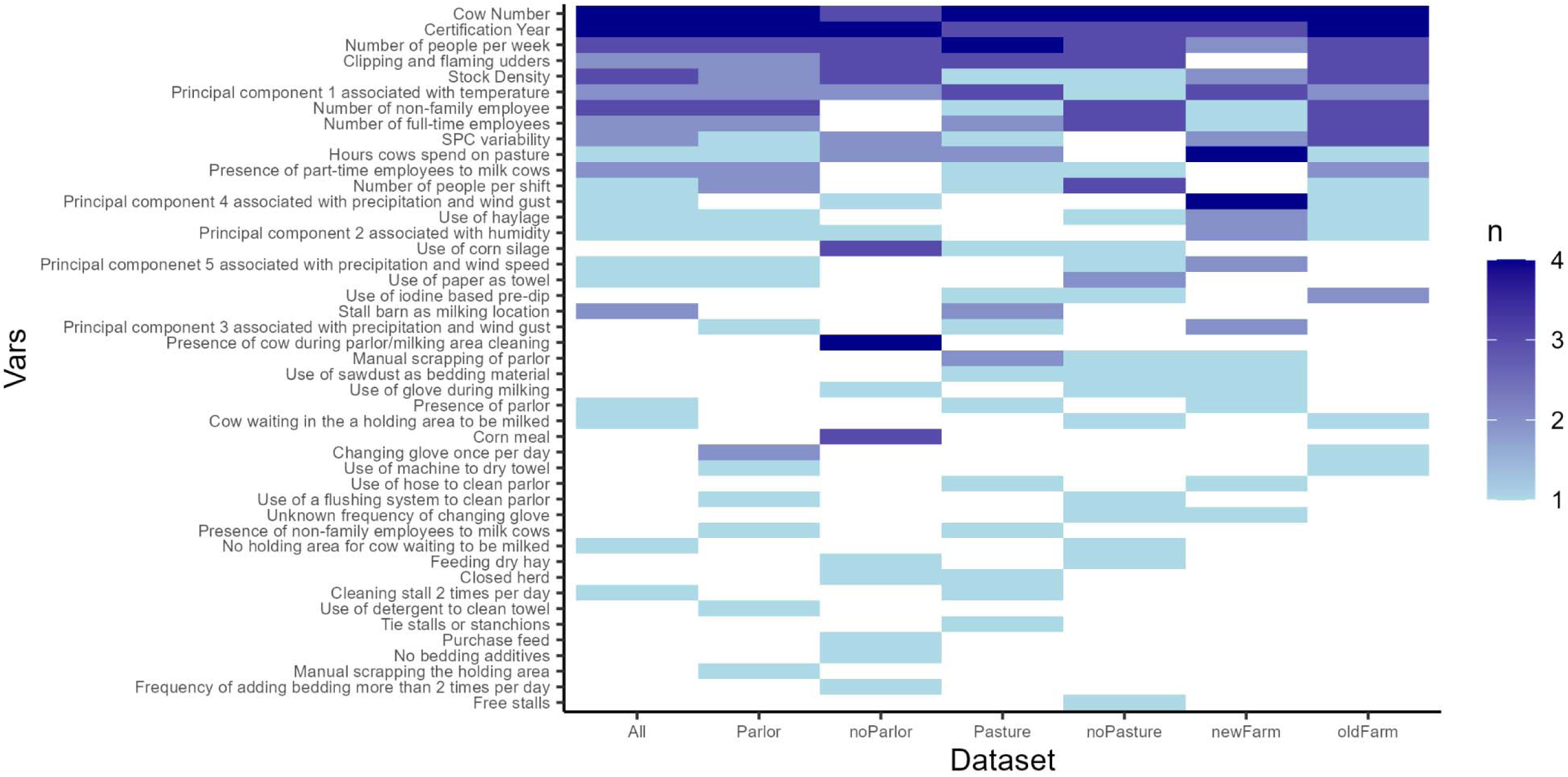
Heatmap for variables identified as top 10 most important from 28 random forest models (4 spore types X 7 data subsets) predicting the spore levels (for mesophilic and thermophilic sporeformers as well as butyric acid bacteria) or presence (for psychrotolerant sporeformers) The seven data subsets include 1 dataset containing all available data, 2 datasets stratified based on whether the farm uses a parlor for milking cows or an alternative approach (e.g., milking in tie stalls), 2 datasets stratified based on the number of years the farm has been certified organic (i.e., whether the farm has been certified for ≤ 9 years or > 9 years), and 2 datasets stratified based on, at each sampling date, whether the lactating cows were exposed to pasture or not. The count value represents the number of spore types that a variable was important for in the model prediction.

From the heatmap, we can visualize that cow number is important for spore levels in organic milk, as it was considered important in 27/28 models. The PDP for cow numbers indicated a trough of spore levels (for BAB, MSC and TSC) at around 100 cows while showed a consistent declining trend for likelihood of detecting spores (for PSC) as cow numbers increased. Considering most farms are not large farms (75% quantile is 300 cows), the trends beyond 300 cows could be biased by a few farms. The second most important variable is the certification year, which suggests the experience of the organic farm has an important role in the spore levels in the milk. We observed in the PDPs that with increasing certification year, the spore levels were expected to decrease, suggesting that farms that have been certified organic for a longer period of time might be more experienced in applying organic practices and in general more consistently applying management practices. Further, the majority of employee-related variables ranked top in the heatmap, including number of different people milking cows per week (identified in the VIP of 21/28 models), number of non-family employees (identified in the VIP of 14/28 models), and number of full-time employees (identified in the VIP of 13/28 models). Analysis of PDPs for the number of different people milking cows throughout a week and number of people per milking shift suggested that the spore levels are the lowest when these two variables are 0, which represents robotic farms. SPC variability, which is included as an indicator of a farm’s ability to manage their milk quality consistently, also demonstrated high importance (identified in the VIP of 11/28 models), except for models constructed for farms without pasture time (e.g., during confinement due to inclement weather). Compared to variables mentioned above, only one farm management practice (i.e., clipping and flaming the udders) was highly ranked. Clipping and flaming the udders was consistently important for all farm styles except for farms certified for at most 9 years, and PDP for this variable consistently indicated that removal of udder hair decreases spore levels or detection likelihood. Among other farm management practices that appear in VIPs multiple times, a few, notably, showed a consistent effect in multiple PDPs. For example, using machines to dry towels and using corn silage consistently decrease spore levels or detection likelihood. In addition, farms that use stalls or tie-stall barns as milking locations as compared to parlors or robots are consistently found to have higher MSC and TSC levels. Conversely, using paper towels to clean udders and using sawdust as bedding consistently increases spore levels or detection likelihood. Some farm management practices, such as type of feed, showed mixed effects depending on the spore type. For example, PDPs for haylage showed that farms using haylage as feed were less likely to detect psychrotolerant sporeformers while having higher levels of mesophilic and thermophilic sporeformers in raw milk. Similarly, using corn silage was shown to decrease the likelihood of detecting psychrotolerant sporeformers but increase the levels of MSC, TSC, and BAB.

In addition to employee-related variables and farm management practices, meteorological factors, which were represented by the principal components, also showed importance in the heatmap. Notably, the principal component 1, which was highly correlated with air temperature (see specific loading plots for principal component extraction at https://github.com/FSL-MQIP/OREI_ML_Inference/tree/main/Figure/PCA) was considered influential in spore levels for all farm styles, supporting that spore levels would likely fluctuate depending on the meteorological conditions and controlling farm practices alone is not guaranteed to maintain consistent milk quality. Therefore, adjustments in management practices may be necessary during certain times of the year when meteorological conditions represent a higher risk of spore contamination.

While it is a simplifying approach to visualize results from VIPs in a heatmap, we need to take precautions in the interpretation as certain variables, even though ranked top 10 in the VIP, might have such low variable importance scores that their effect could be negligible. This is particularly concerning as our model accuracy overall was relatively low as summarized in Table 3.

**Table 3.**
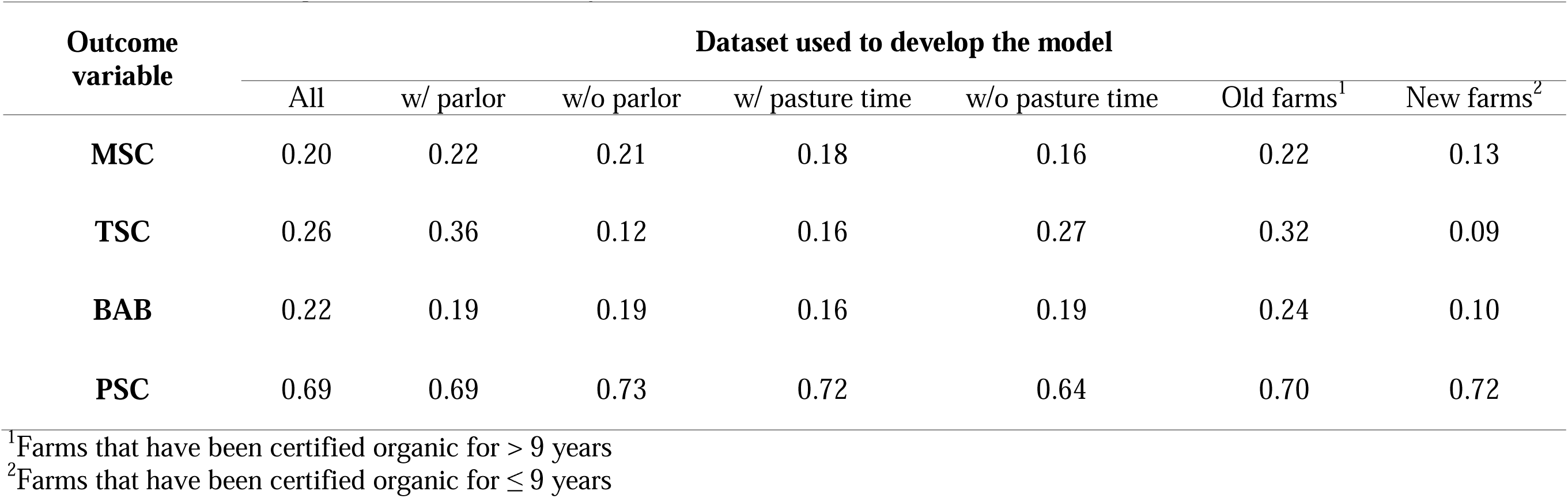
Cross-validation results of random forest regression for predicting log_10_MSC, log_10_TSC, and log_10_BAB (shown in R^2^) and classification for detecting PSC (shown in accuracy).

| Outcome variable | Dataset used to develop the model |  |  |  |  |  | Old farms<sup>1</sup> | New farms<sup>2</sup> |
| --- | --- | --- | --- | --- | --- | --- | --- | --- |
|  | All | w/ parlor | w/o parlor | w/ pasture time | w/o pasture time |  |  |  |
| **MSC** | 0.20 | 0.22 | 0.21 | 0.18 | 0.16 |  | 0.22 | 0.13 |
| **TSC** | 0.26 | 0.36 | 0.12 | 0.16 | 0.27 |  | 0.32 | 0.09 |
| **BAB** | 0.22 | 0.19 | 0.19 | 0.16 | 0.19 |  | 0.24 | 0.10 |
| **PSC** | 0.69 | 0.69 | 0.73 | 0.72 | 0.64 |  | 0.70 | 0.72 |

<sup>1</sup>Farms that have been certified organic for > 9 years
<sup>2</sup>Farms that have been certified organic for $\leq 9$ years

### Data stratification reveals that drivers of organic raw milk spore levels vary by farm practices and characteristics

With the intention to improve the model performance as well as suggest more targeted, individualized spore reduction strategies for farms with different management practices, we stratified our dataset by three factors as previously mentioned, including use of parlor, pasture time, and certification year, respectively. While there are a few overlapping important variables such as certification year, number of cows, etc. (Figure 3), farms with different characteristics varied in other drivers for spore levels. In addition, model performance was also impacted by data stratification, suggesting that certain farms are more sufficiently characterized by the existing data we collected (Table 2) compared to analyzing all farms together in the same model. Notably, we found that in most cases, the model prediction improved for farms that have been certified for more than 9 years, compared to farms that have been certified for at most 9 years. This prediction improvement may indicate that farms that have been certified for fewer years are less consistent in the execution of farm management practices, which could reduce the trustworthiness of survey responses. On the other hand, important variables for farms that have been certified for more than 9 years are mostly concentrated and generally aligned with the overall trend (e.g., cow number, employee-related variables). There are a few farm practices that are more important for farms that have been certified for more than 9 years, but not important for farms that have been certified for at most 9 years, including using iodine-based pre-dip, cows waiting in a holding area to be milked, use of machine to dry towels used during milking preparation, and the use of a hose to clean the parlor. Comparing dairy farms that use and do not use a parlor (e.g., tie stall/stanchion milking), the model performance showed less discrepancy, except for when predicting TSC, in which cases the model performed better predicting the TSC for parlor farms. This potentially suggests that for farms without a parlor, more external factors or contextual information not included in this study can potentially influence the spore levels. The heatmap (Figure 3) showed that farms without parlors were influenced by the use of corn silage, the presence of cows during cleaning the milking area, as well as the use of corn meal whereas glove changing frequency is relatively important on farms not using parlors for milking. Similar to data stratified by parlor, we observed a discrepancy in model performance only in predicting TSC levels when comparing samples that were collected from farms when cows were exposed to pasture versus samples that were collected from farms when cows were not on pasture. It showed that when cows are on pasture, the spore levels are less explainable by only farm practices and meteorological conditions, which could suggest that external factors, such as environmental factors other than meteorological factors are responsible for spore levels when cows are on pasture (e.g., spore levels in the natural environment). Despite that, air temperatures (as represented by principal component 1) and pasture time are still important for 3 (i.e., mesophilic, thermophilic sporeformers and butyric acid bacteria) and 2 (i.e., psychrotolerant sporeformers and butyric acid bacteria) spore types, respectively, when cows are on pasture (Figure 3).

## DISCUSSION

### Spore levels in organic bulk tank raw milk are similar, but slightly higher than those previously reported in conventional bulk tank raw milk

The spore levels reported in our study representing organic dairy farming showed slightly higher results compared to conventional dairy farming, but the difference is relatively small, suggesting that organic dairy farming is not substantially more susceptible to spore contamination. For MSC, we have a positive rate of 94.3% with a mean of 0.76 log_10_CFU/mL and a standard deviation of 0.68 log_10_CFU/mL. The positive rates, mean, and standard deviation summarized in our study are only slightly higher than previously studies, which reported a positive rate of 91% with a mean of 0.50 log_10_CFU/mL and a standard deviation of 0.56 log_10_CFU/mL (Martin et al., 2019) and a positive rate of 85% with a mean of 0.26 log_10_CFU/mL and a standard deviation of 0.56 log_10_CFU/mL (Murphy et al., 2019). Compared to MSC, TSC results showed a lower positive rate of 85.6% with a mean of 0.43 log_10_CFU/mL and a standard deviation of 0.70 log_10_CFU/mL. These values are very similar to data collected from previous studies, in which a positive rate of 87% with a mean of 0.36 log_10_CFU/mL and a standard deviation of 0.51 log_10_CFU/mL (Martin et al., 2019) and a positive rate of 85% with a mean of 0.26 log_10_CFU/mL and a standard deviation of 0.52 log_10_CFU/mL (Murphy et al., 2019) were reported. Similar to TSC, BAB data from our study are not substantially different from previous studies (Vissers et al., 2007a; Shi et al., 2023). Organic milk samples evaluated in this study had a 94.4% positive rate with a mean of 2.19 log_10_MPN/L and a standard deviation of 0.65 log_10_MPN/L. By comparison, Vissers et al. (2007a) reported a mean of 2.10 log_10_MPN/L with a standard deviation of 0.71 log_10_MPN/L while Shi et al. (2023) reported a positive rate of 87% with a mean 1.79 log_10_MPN/mL and a standard deviation of 0.59 log_10_MPN/L for 12 NY farms enrolled in the study. Similar to the other spore levels reported here, the levels and variance in psychrotolerant spore count found in this study were in line with what research on conventional raw milk has reported. In our study, 72.1% of samples tested positive for PSC. The mean (-1.50 log_10_MPN/mL) and standard deviation (0.68 log_10_MPN/mL) are both in a similar scale compared to a previous study which reported the PSC in selected NY farms to follow a normal distribution with a mean of -0.72 log_10_MPN/mL and a standard deviation of 0.99 log_10_MPN/mL (Masiello et al., 2014). Overall, the prevalence and concentration distribution for sporeformers in raw milk collected from organic dairy farms are comparable to results previously reported on the conventional farms. While our study did not investigate the similarities in microbial diversity, one previous study (Coorevits et al., 2008) reported that aerobic sporeformer flora were highly similar between milk samples collected from organic and conventional dairy farms.

Although it is not within the scope of this study, using microbial data generated from this study with characterized between-farm and within farm variability will be valuable for incorporating variability across different geographical locations and seasons. While previous studies characterized the spore levels in conventional farms, our dataset can be used as input for 2-dimensional Monte Carlo simulation models, which have been increasingly used to predict microbial food spoilage (Nielsen et al., 2021; Lau et al., 2022; Qian et al., 2022, 2023; Snyder et al., 2024), tailored specific to organic farms. More importantly, dairy farmers and processors can further subset out this dataset to farms that share similarities (e.g., similar climate and management practices) to their own conditions to improve model precision. These models can provide additional digital tools for organic farmers who want to utilize their farm data for predicting the dairy spoilage in the entire supply chain.

### Reducing spore levels in organic bulk tank raw milk will require an individualized, targeted approach that accommodates farm characteristics, time of year, and meteorological factors

We identified several farm practice factors that are influential to spore levels as shown in Figure 3. The importance of these factors has been previously investigated by other studies. For instance, the most important variable in our study, the number of cows, which represents the herd size, was also previously found to be associated with levels of sporeforming bacteria in bulk tank milk (Masiello et al., 2014; Miller et al., 2015). Specifically, one study concluded that farms with less than 200 cows are more likely to have higher psychrotolerant spore levels (Masiello et al., 2014) while in a different study, large herd size was associated with lower mesophilic and thermophilic spore levels (Miller et al., 2015). In addition to spore levels, large herd size was also found to correlate with lower SPC and SCC, which are indictors of raw milk quality (Ingham et al., 2011). However, our PDP for the herd size suggests that the effect of this variable might not be linear and monotonic as previously reported. When other conditions were held constant, increasing herd size initially decreases levels of mesophilic and thermophilic sporeformers and butyric acid bacteria and then increases these spore levels until the effect reaches a nearly non-substantial level when the herd size is above approximately 200 cows. However, the PDP for PSC showed that increasing the herd size always decreases the likelihood of detecting psychrotolerant sporeformers. As 75% of our farms have less than 300 cows, any predicted effect of herd size beyond 1000 cows in the PDP might not be representative and therefore should be interpreted with caution. In addition, as herd size is a compound variable (associated with other management factors such as financial capacity), additional work is needed to disintegrate this variable into representative variables that biologically drive spore levels. Following the herd size, the number of years an organic farm has been certified was also important. PDPs for this variable consistently showed a decrease in spore levels when the farm has been certified organic for a longer period. Practically, this suggests that organic farms that have been certified for a longer period of time might have more experience or more control over their milk production. While no previous study has evaluated the effect of certification year on the spore levels, other studies (Evanowski et al., 2020, 2023) did, however, support that additional training for milking staff on the milking preparation was an effective intervention strategy to reduce spore levels, which may be a factor that is influencing this result.

After the herd size, employee-related variables, such as number of people per week, number of non-family employees, and number of full-time employees, appeared frequently in the VIPs (Figure 3). Similar to the herd size, these variables are most likely proxy for other factors (e.g., fewer full-time employees might indicate more employees are less well-trained and prone to behave undesirably during milking). Close examination of the PDPs for these variables revealed that while effects for number of full-time employees and number of non-family employees are too negligible to draw any meaningful conclusions, when number of people per week is 0, which indicates automated milking system (AMS), the predicted mesophilic and thermophilic spore levels are at the lowest. Among the samples collected from farms with AMS, we observed decreased positive rates for mesophilic (64.7% compared to 94.3% for all samples) and thermophilic sporeformers (45.0% compared to 85.6% for all samples). However, since we only enrolled 4 robotic farms in our study, the generalizability of this finding needs further investigation. As supported by a published review paper (Hogenboom et al., 2019), contradictory findings were reported for the effect of AMS on the microbiological quality of milk, depending on the correct implementation of milking hygiene practices in AMS. This, alongside our findings, suggests that AMS has the potential to reduce the spore levels in milk if consistency in automation of milking hygiene can be achieved.

Among principal components that represent meteorological factors, principal component 1, which is highly correlated with air temperature, is one of the most important variables influencing the spore level. A previous study (Vissers et al., 2007b) showed that *B. cereus* spore levels are higher by 0.5 ± 0.02 log_10_ spores/L in the summer compared to other times, presumably due to increased growth at higher temperatures, although this difference was not statistically evaluated. A different study (Martin et al., 2019) applied multimodel inference to evaluate the effect of average temperature and concluded that this variable had a negligible impact compared to spore levels in the environment as well as various farm practices. The importance of temperature in our study might be attributable to organic farming practices, which require pasture time for cows when weather conditions permit.

Compared to farm size, certification year, employee-related factors, and temperature, farm practices were less consistently important in the heatmap. Among the farm practices, two categories showed frequent occurrence in the heat map; these two categories are (i) udder and teat hygiene, which includes predictor variables clipping and flaming, and use paper as towel, use of iodine-based pre-dip, use of a machine to dry towels, and use of detergent to sanitize towels, and (ii) bedding, which includes variables such as sawdust bedding and frequency of topping up bedding. The importance of these two categories of farm practices is consistent with previously published literature. For example, our results consistently identified that removal of udder hair was associated with lower spore levels, consistent with what Martin et al. (2019) reported, specifically that udder hygiene conditions were found important to MSC, TSC, and BAB while udder clipping and flaming was shown important to specially thermoresistant spore enumeration and BAB (Martin et al., 2019). Meanwhile, practices used to improve udder hygiene, such as by improving laundered towel preparation using detergent and drying was shown to reduce mesophilic spore levels by 37% and thermophilic spore levels by 40% in an intervention study (Evanowski et al., 2020). Our PDPs showed that machine drying towels used for milking preparation was predicted to reduce MSC in farms that had been certified for longer than 9 years by approximately 0.1 log_10_ CFU/mL and reduce the likelihood of detecting psychrotolerant sporeformers in farms using a parlor for milking by approximately 10 percentage points. In addition to the udder and teat hygiene, the choice of bedding and the frequency of adding bedding were both reported to correlate with spore levels in bulk tank milk. For example, sawdust bedding was shown to be correlated with lower MSC while straw bedding was shown to be correlated with lower TSC (Miller et al., 2015). However, a different study (Murphy et al., 2019) showed that organic beddings, such as sawdust, correlate directly with higher spore levels in unused and used beddings and might indirectly correlate with spores in bulk tank milk. PDPs for beddings in our study showed negligible difference in spore levels for different bedding types overall, except that for organic dairy farms that have been certified for at most 9 years, using sawdust is 10 percentage points more likely to detect psychrotolerant sporeformers. The seemingly conflicted findings on the effect of bedding materials suggest material type alone is only an intermediate variable and that monitoring the spore levels in bedding as a direct indicator as contamination source is important for predicting the spore levels in the raw milk.

Although several farm practice variables ranked among top in multiple VIPs generated from models constructed using different subsets of data, it is difficult to interpret the reason that certain variables only showed importance for particular farm styles (e.g., presence of cow during milking area cleaning is predominantly important for farms without parlor). Given that a variable can have a low variable importance score and still rank top 10 in a model that has poor performance, whether these sparsely important variables are drivers for spore levels should be interpreted with caution, and therefore it is important to conduct further farm experiments to validate the hypothesis from these findings. In addition, as most variables only provide minimal effects to the spore levels in the PDPs, it is important to evaluate the synergistic effects of multiple variables simultaneously to understand the risks of spore levels in raw milk. While it would be valuable to demonstrate this via multiple dimensional partial dependence plots, the number of combinations would be impractical to comprehensively exhibit here. Conceptually, our model can be practically used to interpret, for example, risks of BAB levels are expected to be higher in an organic dairy farm that is certified for at most 9 years, does not perform udder clipping and flaming, and has limited pasture time for cows when the weather conditions do not allow cows on pasture.

### Integration of novel data streams with state-of-art machine learning algorithms is potential to enable real-time monitoring of spore levels in the future

In this study we used a machine learning approach to examine associations between farm management survey data and spore levels in organic bulk tank raw milk samples. Surveys have been frequently used in the past (Masiello et al., 2014, 2017; Miller et al., 2015; Murphy et al., 2019) to collect information from dairy farmers regarding farm management as it is a common tool in epidemiological studies (Dohoo et al., 2003b) to study the relationship between the exposure and outcome, which in our case, are farm practices and spore levels in milk, respectively. While it is convenient to use surveys for cross-sectional studies, it may suffer from recall bias (Dohoo et al., 2003c), which refers to the situation where respondents may not accurately report their behaviors. In addition, a survey tends to collect aggregate information over a period, which is appropriate to obtain an overview for farm operations but will inevitably neglect execution-level details. In the context of dairy farm operations, these details can lead to high uncertainty in raw milk quality as farms, especially those with more heterogeneity in employee composition and rotation and without formal employee training, can have rare events that differ from survey responses (e.g., a part-time employee who does not change gloves as frequently as the farm manager reported). Therefore, while surveys have been proved to be a suitable tool to represent overall farm practices and investigate effect of management-level factors, it falls short as a data source for developing a model capable of real-time predictions of spore levels. This limitation of surveys can be overcome by novel data streams such as video surveillance data. With recent advancements in computer vision for agriculture automation (Tian et al., 2020), cameras have been more frequently suggested as a potential equipment for establishing an intelligent dairy farm system for cattle management (Niall et al., 2019; Zhang et al., 2023). As the udder is the key entry point for spore contamination, monitoring the cleanliness of teats and udders would be critical. Previous studies have proven that using these computer vision tools were sufficiently precise to predict milk yield based on udder traits (Shorten, 2021) as well as detect teats for application of teat cup attachment (Lu et al., 2021). It is reasonable to assume that these tools can be extended to monitor the teat during the milking process and detect any anomaly behaviors from employees that increase the risk of spore contamination. In addition to failure to adequately clean the udder, cleaning the parlor or housing area when cows are present might also increase the likelihood of spore contamination, as spores from soil and manure, which have been shown to have very high concentrations of spores (Martin et al., 2019), could be spread onto the cows when parlor or housing area is being cleaned with hoses that yield large pressure. This, again, is another example of undesirable behavior that establishing a computer vision system in the parlor or housing area can monitor. This computer vision system at the housing area could also be used to monitor the frequency of adding beddings, which is associated with PSC, MSC, and TSC (Martin et al., 2019).

While video surveillance is appropriate for objectively recording animal and human behaviors, recording verbal responses, in the form of interview, from milking staff or farm managers can provide essential information of farm management, sometimes qualitative, such as the type of pre-dip used. Recently, large language model (LLM) is rising as revolutionizing tool that can be fine-tuned to downstream tasks specific to human language (Naveed et al., 2023). This type of model can analyze unstructured data such as interview transcripts, which can, compared to surveys, provide (i) more contextual information that represents heterogeneous farm characteristics and (ii) anecdotes that occur rarely but could be relevant directly or indirectly to spore levels transmitted to milk. For example, a manager might mention that the farm implements a sprinkler system to cool off cows on hot summer days, which can contribute to increased contamination of the udder by washing down the dirt to the teats. LLMs are not only a promising tool for interpreting survey responses but also show potential in automating the interview process as a conversational tool that can chat with managers in real-time (Sahijwani et al., 2023), which would allow collecting responses from dairy farms more regularly with less labor and time commitment as well as recording rare events promptly. Currently, while our machine learning approach using survey and meteorological data was insufficient for accurately predicting spore levels in organic raw milk, it is possible to envision the development of an end-to-end application that combines (i) incidence of undesirable behaviors extracted from on-farm video data, (ii) insights extracted from LLM conversational tool, and (iii) meteorological data reported from nearby weather stations for enabling real-time risk prediction for spore levels in raw milk.

## CONCLUSION

Longitudinal data on spore levels in organic raw milk from our study can be incorporated into an existing spoilage risk assessment model to evaluate the risk of finished product spoilage based on initial spore levels. The machine learning approach in this study identified important farm practices for various farm styles. The developed models can practically (i) generate hypotheses to guide future intervention studies in choosing potential strategies to evaluate and (ii) estimate spore levels or detection likelihood for individual farms with varying management practices under different meteorological conditions. These models serve as an initial step towards incorporating other novel data streams, such as video surveillance and automated verbal responses, to establish a real-time monitoring tool for key quality parameters of organic raw milk

## ACKNOWLEDGEMENTS

This study was funded by United States Department of Agriculture (Washington, DC) Organic Research and Extension Initiative (USDA OREI) grant no. 2019-51300-30242. The authors acknowledge that N. Martin is a section editor for the Journal of Dairy Science and have not stated any conflicts of interest.

